# Functional characterization of germline variants in the *TMEM127* tumor suppressor reveals novel insights into its membrane topology and trafficking

**DOI:** 10.1101/2020.04.08.031039

**Authors:** Shahida K. Flores, Yilun Deng, Ziming Cheng, Xingyu Zhang, Sifan Tao, Afaf Saliba, Irene Chu, Exing Wang, Ricardo C. T. Aguiar, Patricia L. M. Dahia

## Abstract

**Purpose:** To better understand the function of the transmembrane protein TMEM127, a poorly known tumor suppressor gene associated with pheochromocytomas, paragangliomas and renal carcinomas, we evaluated patient-derived germline variants.

**Methods:** Subcellular localization and steady-state levels of 21 tumor-associated, transiently expressed *TMEM127* variants were compared to the wild-type protein using immunofluorescence and immunoblot analysis, respectively, in cells genetically modified to lack endogenous TMEM127. Membrane topology and endocytic mechanisms were also assessed.

**Results:** We identified three subgroups of mutations and determined that 15 of the 21 variants (71%), including 9 of 15 missense variants (60%), are pathogenic or likely pathogenic, through loss of membrane binding ability, stability and/or internalization capability. Investigation into an N-terminal cluster of missense variants uncovered a previously unrecognized transmembrane domain, indicating that TMEM127 is a four-, not a three-, transmembrane domain-containing protein. Additionally, a C-terminal variant with predominant plasma membrane localization revealed an atypical, extended acidic, dileucine-based motif required for TMEM127 internalization through clathrin-mediated endocytosis.

**Conclusion:** We characterized the functional deficits of several germline *TMEM127* variants and identified novel structure-function features of TMEM127, namely, a fourth transmembrane domain and an endocytic motif. These findings will assist in *TMEM127* variant interpretation and will help guide future studies investigating the cellular role of TMEM127.

---

Pheochromocytomas (PHEOs) and paragangliomas (PGLs) are rare, neural crest-derived, catecholamine-secreting tumors that arise in the adrenal medulla and along the paraganglia, respectively^1,2^. These tumors are considered genetically heterogeneous and highly heritable with 12-20 susceptibility genes, with varying degrees of validation, identified to date^1-3^. Additionally, PHEOs and PGLs frequently co-occur with other tumors, including renal cell carcinomas (RCCs), in well-established hereditary tumor syndromes, indicating a shared genetic predisposition^4,5^.

The transmembrane protein-encoding gene *TMEM127* has been identified as a susceptibility gene in PHEOs^6-9^. More rarely, variants have also been detected in PGLs and RCCs^10-13^. Germline variants in *TMEM127* account for approximately 2% of all cases of PHEOs^9,14^. The *TMEM127* gene encodes a poorly characterized but highly conserved, ubiquitously expressed 238-amino acid protein with no significant homology to other proteins and no apparent functional domains other than three predicted transmembrane domains^6^. TMEM127 has been characterized as a tumor suppressor, a negative regulator of mTOR signaling and an endo-lysosomal membrane protein^6,10,11,15^. Mutant *TMEM127* PHEOs and PGLs display an expression profiling related to ‘cluster 2’, a group that comprises other mutant tumors predominantly associated with increased kinase signaling^6^. Wild-type (WT) TMEM127 colocalizes with markers of the plasma membrane, but more extensively with various endomembrane structures (early endosome, late endosome and lysosome), displaying a characteristic predominantly punctate appearance^6,10,15^. Altered subcellular localization, specifically a diffuse cytoplasmic distribution, was previously detected in patient-derived *TMEM127* variants that disrupt transmembrane domains^6,7,11^. In addition, TMEM127 coprecipitates with proteins involved in amino acid-mediated mTOR recruitment to the lysosome and, by still poorly defined mechanisms, contributes to reduced mTORC1 signaling^15^. Furthermore, recent *in vivo* studies have also indicated a role for TMEM127 in nutrient sensing, glucose and insulin homeostasis and mTORC2 signaling^16^. However, the precise physiological role of TMEM127, and its mechanisms of action and regulation, remain poorly defined.

Given the limited information on the structure, function and regulation of TMEM127, ascertaining the pathogenicity of germline *TMEM127* variants has been a major challenge. As a recognized tumor suppressor gene with Tier 1 status in the COSMIC Cancer Gene Census (COSMIC v90) and as a PHEO/PGL/RCC susceptibility gene, *TMEM127* has been included in several commercially available and research-based hereditary cancer genetic screening panels^17^. Over one hundred unique, low frequency, germline *TMEM127* variants have been reported in the context of familial or sporadic PHEOs, PGLs and/or RCCs^4,7-10,12-14,18-27^. However, many of these *TMEM127* variants, especially missense changes, are currently classified as variants of uncertain significance (VUS). As the American College of Medical Genetics and Genomics and Association for Molecular Pathology (ACMG-AMP) guidelines recommended that a VUS not be used for clinical decision making, efforts to reclassify these variants are needed^28^.

In the present study, in an effort to improve our understanding of TMEM127 function and thus predict pathogenicity of variants detected in clinical testing, we systematically evaluated 21 tumor-associated, germline *TMEM127* variants by subcellular localization and steady-state level analysis. We found that at least 15 of these variants, including 9 of 15 missense variants, can be classified as pathogenic or likely pathogenic. Importantly, these analyses provided the backdrop to identify and characterize previously unrecognized structure-function features of TMEM127 that shed light on TMEM127 function.

## MATERIALS AND METHODS

### Patient-derived *TMEM127* Variants

We generated a list of germline *TMEM127* variants for analysis based on published literature, a publicly available database (ClinVar), as well as unpublished variants from our own cohort (obtained through an ongoing study, NCT03160274). Clinical features related to those variants will be reported in a separate manuscript.

### Plasmids, Cell Culture and Transfections

Site-directed mutagenesis using Phusion High-Fidelity DNA Polymerase (Thermo Fisher) and complementary primer pairs (sequences available on request) was used to generate TMEM127 variants in the pEGFP-C2 (Clontech Laboratories) and pCMV6-XL5 (Origene) constructs containing the TMEM127 wild-type (WT) coding sequence (NM_017849.3) as previously described^6^. An MSCV construct which generates TMEM127-WT fused to a Flag tag at its C-terminal (TMEM127-Flag) combined with a biscistronic GFP reporter was also used. These constructs were transiently transfected into HEK293FT cells (ATCC) CRISPR/Cas9-edited to knockout TMEM127 (TMEM127 KO) as we previously reported^11^. Transfections were performed using Lipofectamine 2000 (Invitrogen) according to the manufacturer’s recommendations. HEK293FT cells stably expressing GFP-TMEM127 WT were also generated using standard methods. These cell lines were maintained in Dulbecco’s Modified Eagle’s Medium (DMEM) supplemented with 10% fetal bovine serum (FBS) and 1% penicillin-streptomycin.

### Immunofluorescence microscopy

HEK293FT TMEM127 KO cells transfected with the GFP-tagged TMEM127 WT and variant constructs were split 8-10 hours post-transfection and plated onto coated glass coverslips. At 48 hours post-transfection, the cells were washed with PBS, fixed with 4% paraformaldehyde for 15-20 minutes, washed and stained with 4′,6-diamidino-2-phenylindole (DAPI) for nuclear signal, then mounted onto slides using VectaShield Anti-fade Mounting Medium (Vector Labs). Additional steps when the untagged TMEM127 construct was used or for colocalization analysis included: blocking and permeabilization after fixation with 5% Horse Serum and 0.1% Triton X-100 in PBS for 1 hour, then overnight probing at 4°C with primary antibodies for either TMEM127 (1:500, rabbit, Bethyl Labs), plasma membrane marker Na^+^K^+^-ATPase (1:200, rabbit, CST), lysosomal marker LAMP1 (1:200, mouse, CST) and/or early endosomal marker EEA1 (1:100, rabbit, CST) followed by 1 hour probing at room temperature with secondary antibody (either AlexaFluor-568 at 1:1200 or Cy5 at 1:500 for rabbit; Mouse Texas Red at 1:500 for mouse) before DAPI staining and mounting. Fluorescence images were captured with a Zeiss LSM710 Confocal Laser Scanning Microscope (Carl Zeiss) using the 63X oil objective. The 405nm laser was used to collect blue (DAPI) signal, the 488nm laser for green (GFP) signal, the 561nm laser for red signal (AlexaFluor-568 or Mouse Texas Red) and the 594nm laser for far-red signal (Cy5)^15^. Confocal images are representative of at least 25, but usually more than 50 transfected cells examined in two to five independent experiments.

### Immunoblot analysis

Transiently transfected HEK293FT TMEM127 KO cells collected 24, 48 and 72 hours post-transfection were lysed with a buffer containing NP-40 detergent plus Halt Protease and Phosphatase Inhibitor (Thermo Fisher). Whole protein lysates (30ug) were loaded onto a 12% acrylamide gel after boiling with denaturing loading buffer. After electrophoresis, proteins were transferred unto a PVDF membrane which was probed using antibodies for TMEM127 (1:10,000, rabbit, Bethyl Labs), GFP (1:2000, rabbit, CST), and B-actin (1:15,000, mouse, CST). Blots were developed with chemiluminescent detection.

### *In Silico* Analysis

We used the bioinformatics tool PolyPhen-2 (http://genetics.bwh.harvard.edu/pph2/) to predict the impact of amino acid substitutions resulting from missense *TMEM127* variants. To estimate the location of TMEM127 transmembrane domains, the 238-amino acid sequence of TMEM127 was queried across several protein prediction algorithms: TMHMM (www.cbs.dtu.dk/services/TMHMM/)^29^, UniProt (www.uniprot.org), PolyPhobius (www.phobius.sbc.su.se/poly.html), PHYRE 2 (www.sbg.bio.ic.ac.uk/phyre2/), SOSUI (http://harrier.nagahama-i-bio.ac.jp/sosui/)^30^,Philius (www.yeastrc.org/philius/pages/philius/runPhilius.jsp;)^31^,TOPCONS (http://topcons.cbr.su.se/pred/) and TMPred (https://embnet.vital-it.ch/software/TMPRED_form.html), using the default settings for each program. Several of these programs also generate a prediction for terminal orientation and/or localization of the protein. For generation of 2D models for TMEM127, the visualization tool Protter (www.wlab.ethz.ch/protter/start/)^32^ was used.

### Antibody Accessibility Assay

HEK293FT TMEM127-KO cells were transiently transfected with the MSCV driven vector expressing C-terminal, Flag tagged TMEM127 WT (TMEM127-Flag) with a bicistronic GFP reporter^6^ and plated onto coverslips as described above. After fixation, cells were incubated for 15-20 minutes in PBS containing either: a) 5% horse serum only (‘no detergent, no permeabilization’), b) 5% horse serum plus 0.01% digitonin (‘selective permeabilization’), or, c) 5% horse serum plus 0.1% Triton X-100 (‘full permeabilization’). Next, cells were incubated in primary antibodies targeting TMEM127 on the N-terminal (Bethyl Laboratories, rabbit, 1:500) and Flag on the C-terminal (Cell Signaling Technologies, mouse, 1:500) for 2 hours at room temperature. After PBS washes, cells were incubated in secondary antibodies Mouse Texas Red X (ThermoFisher, anti-mouse, 1:500) and Cy5 (anti-rabbit,1:500), for 1 hour at room temperature. Cells were then prepared for confocal microscopy as described above.

### Clathrin and caveolin knockdown studies

For clathrin knockdown, HEK293FT cells stably expressing GFP-TMEM127-WT were either transduced with lentiviral particles expressing pLKO.1 shRNA constructs targeting against clathrin heavy chain (CLTC81and CLTC82), a gift from Dr. Lois Mulligan^33^, or scrambled shRNA. Experiments used combination of both CLTC81 and CLTC82 supernatants. Lentiviral supernatant generation was processed as reported^6^. Caveolin knockdown was achieved by transfection with siRNA against caveolin (MISSION® esiRNA human CAV1, Sigma) or negative control siRNA (MISSION® siRNA Universal Negative Control #1, Sigma), respectively, and processed using standard methods^6^. Cells were collected 48h or 72h after transduction or transfection, respectively, for confocal microscopy. An aliquot of cells was also collected to verify knockdown efficiency by immunoblot analysis using clathrin heavy chain (CLTC) and caveolin (CAV1) antibodies (both CST, rabbit, 1:1000). Fixed cells were stained with the plasma membrane marker Na^+^K^+^-ATPase (Cell Signaling Technology, rabbit, 1:200) and imaged by immunofluorescence microscopy. Experiments were performed in biological triplicates.

### Data Collection and Analysis

All transfections, transductions and confocal microscopy experiments were performed at least twice, but in most cases, three times. In total, there were at least 25 transfected cells with interpretable results (i.e., non-dying, non-saturated fluorescence) per variant and/or condition. The ImageJ software program was used to convert raw imaging files into JPEG images for visualization of subcellular localization. For confocal images where colocalization was investigated, the Colocalization Threshold feature of Fiji/ImageJ was used to determine the Pearson’s Correlation Efficient for a subset of cells from each variant or condition, as we reported^15^. ImageJ-generated JPEG images were also used for semi-quantitative evaluation of band intensity from immunoblot scans to estimate steady-state levels of mutant TMEM127 variants over the specified time course relative to a loading control and to the WT construct. GraphPad Prism was used to generate scatter plots. A Student’s t-test or one-way ANOVA (for more than 2 comparison groups) was used to determine statistical significance. Differences were considered statistically significant when p<0.05.

## RESULTS

### Subcellular localization analysis reveals three types of tumor-associated, germline *TMEM127* variant distribution

We selected 21 tumor-associated, germline *TMEM127* variants (15 missense variants, 3 frameshift variants, 1 N-terminal truncating variant, 1 in-frame insertion variant and 1 in-frame deletion variant) for functional characterization (Table 1). These variants were chosen to represent tumor-associated *TMEM127* variants from across the 238-amino acid TMEM127 protein. First, we observed the subcellular localization of GFP-tagged TMEM127 WT and variant proteins transiently expressed in HEK293FT TMEM127-KO cells at 48 hours post-transfection. The localization pattern of each variant was compared to WT, which we found to display a predominantly punctate appearance, which we previously showed to co-localize to various endosomal domains and the lysosome^6^, but also with approximately 20% of signals detectable at the plasma membrane under steady state culture conditions (Fig1a, 1b). We found that all TMEM127 variant proteins separated distinctly into three subcellular localization patterns (Fig. 1c, 1d). Of the 21 variants evaluated, 8 were punctate/endomembrane (similar to WT), 12 were diffuse/cytoplasmic and 1 was predominantly plasma membrane bound (Fig. 1b, Table 1). To ensure that our observations were not an artifact of the GFP tag, representative variants from each of these three categories were also independently examined using untagged TMEM127 constructs and showed identical patterns (Supplementary Fig. S1a).

**Table 1.**
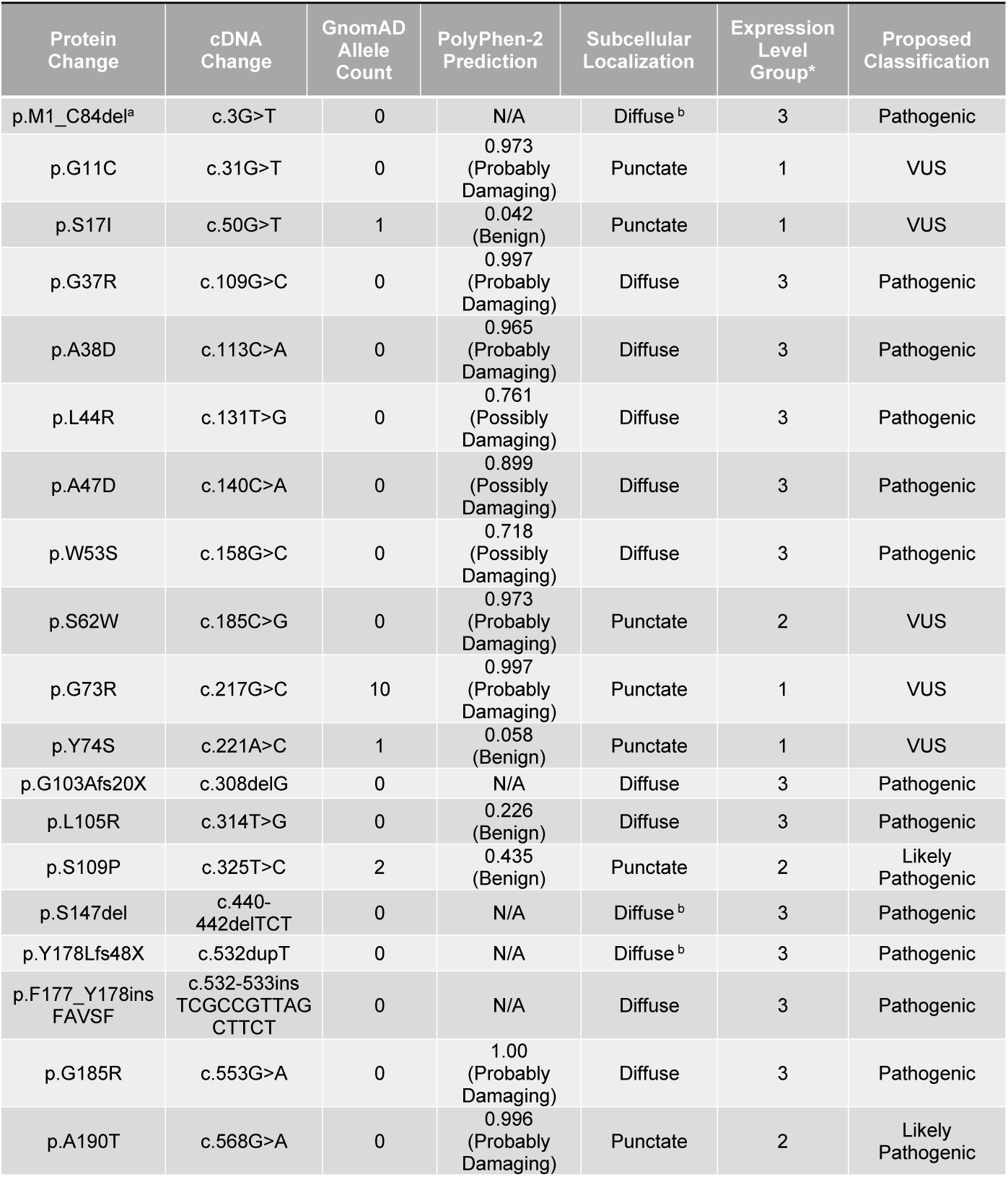
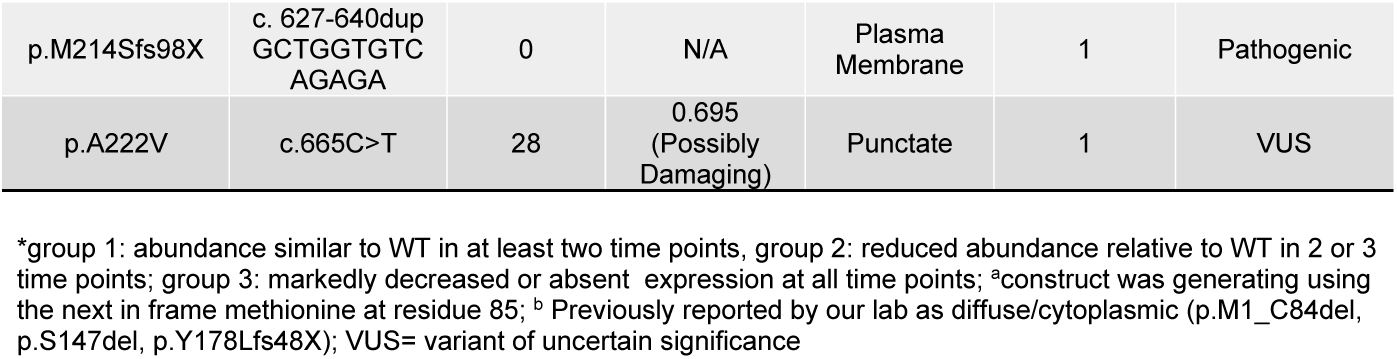
Features and Functional Characterization of Tumor Associated, Germline *TMEM127* Variants

**Fig. 1.**
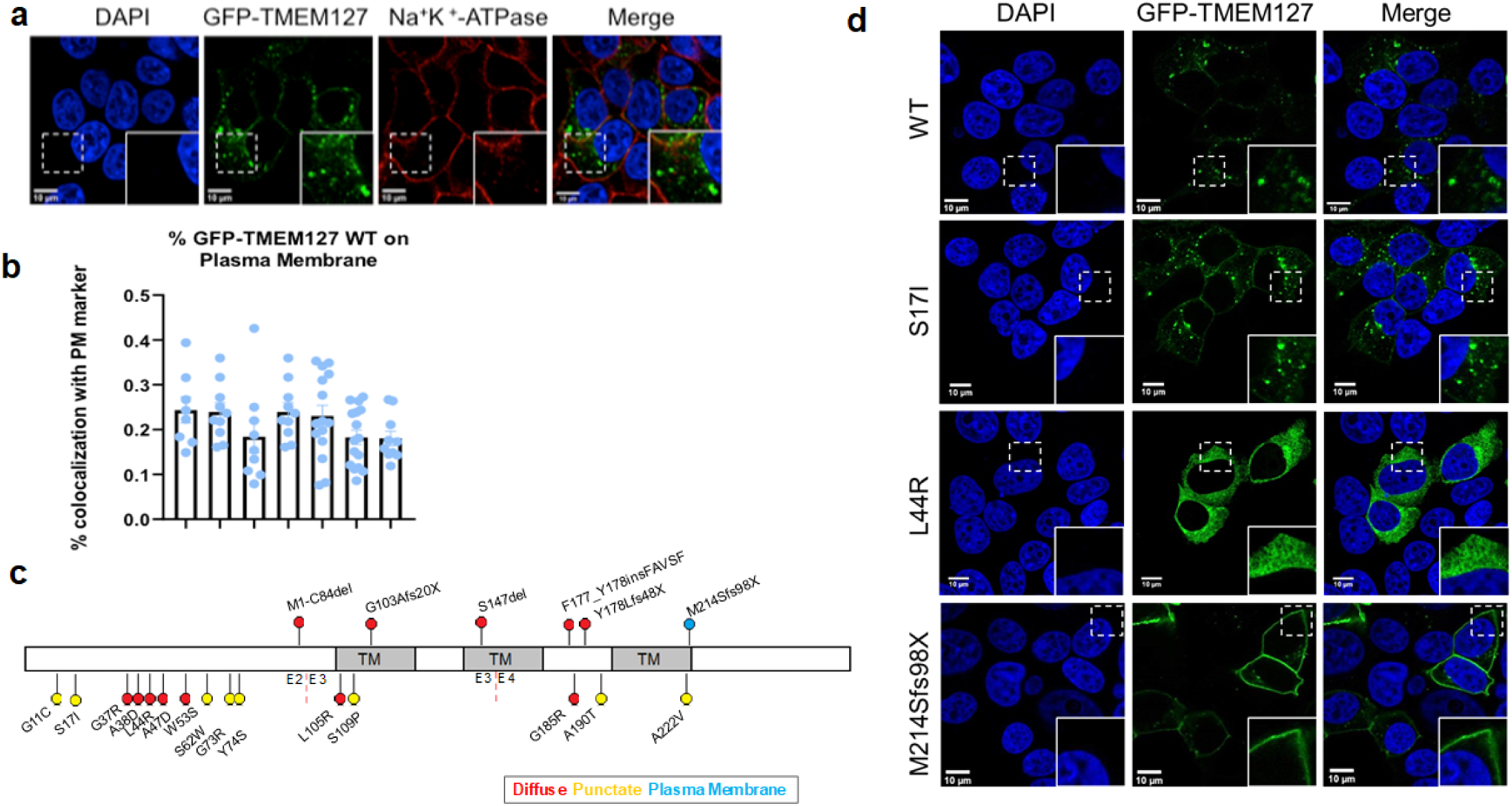
Subcellular localization of tumor-associated germline TMEM127 variants. (a) Representative images from immunofluorescence analysis of HEK293FT TMEM127-knockout (KO) cells transiently expressing GFP-tagged TMEM127 WT (green) showing predominant punctate (endomembrane) appearance, but also plasma membrane distribution, by colocalization with a plasma membrane marker NA^+^K^+^ATPase (red); (b) Quantification of plasma membrane colocalization of GFP-TMEM127 transfected in HEK293FT TMEM127 KO cells from seven independent experiments and 78 cells (green channel) indicating that 21.4% (SD+-0.07) of TMEM127 is detected at the cell surface; (c) Graphic representation of the location of the 21 *TMEM127* variants examined and their corresponding subcellular localization with frameshift, truncating, in-frame insertion and in-frame deletion variants represented by upward facing lollipops and missense variants by downward facing lollipops (punctate/endomembrane = yellow, diffuse/cytoplasmic = red, plasma membrane = blue); (d) Representative images from immunofluorescence analysis of HEK293FT TMEM127-KO cells transiently expressing the indicated variants separated distinctly into three subcellular localization patterns: punctate/endomembrane (represented by p.S17I), diffuse/cytoplasmic (represented by p.L44R) and predominantly plasma membrane (represented by p.M214Sfs98X). WT-TMEM127 is shown as reference. Nuclear staining with DAPI (blue channel) is also shown. Table 1 displays the full list of variants tested.w

We then assessed the localization pattern of each variant, the type of sequence variation and the corresponding position along the TMEM127 amino acid sequence relative to the three previously reported transmembrane domains. In our earlier studies, we have observed that variants that disrupt one or more transmembrane domains typically show a diffuse, cytoplasmic distribution^7,10,11^. All of the eight variants that were punctate were missense substitutions, of which six (p.G11C, p.S17I, p.S62W, p.G73R, p.Y74S, p.A222V) were located outside of transmembrane domains and only two (p.S109P, p.A190T) were situated within transmembrane domains (Fig. 1c, 1d, Suppl Fig1a). The other seven missense variants were diffuse/cytoplasmic with two variants (p.L105R, p.G185R) residing within transmembrane domains and five variants (p.G37R, p.A38D, p.L44R, p.A47D, p.W53S) located in an N-terminal region not previously known to have any recognizable domains (Fig. 1c, Table 1). The in-frame insertion and in-frame deletion variants (p.F177_Y178insFAVSF, p.S147del) and two frameshift variants (p.G103Afs20X, p.Y178Lfs48X), which were diffuse, disrupted one or more transmembrane domains (Fig. 1c, Supplementary Fig. S1b, Table 1). The N-terminal truncating variant (p.M1_C84del), which we previously described^7^, despite not disrupting a reported transmembrane domain was also diffuse in the cytoplasm (Fig. 1c, Fig.3b, Supplementary Fig. S1b), as detailed below. Lastly, the remaining frameshift variant (p.M214Sfs98X), located downstream of the last transmembrane domain, hence maintaining the integrity of all transmembrane domains, was predominantly localized to the plasma membrane, a distinctly unique pattern from the other variants (Fig. 1c, 1d; Supplementary Fig. S1a, 1b).

**Fig. 2.**
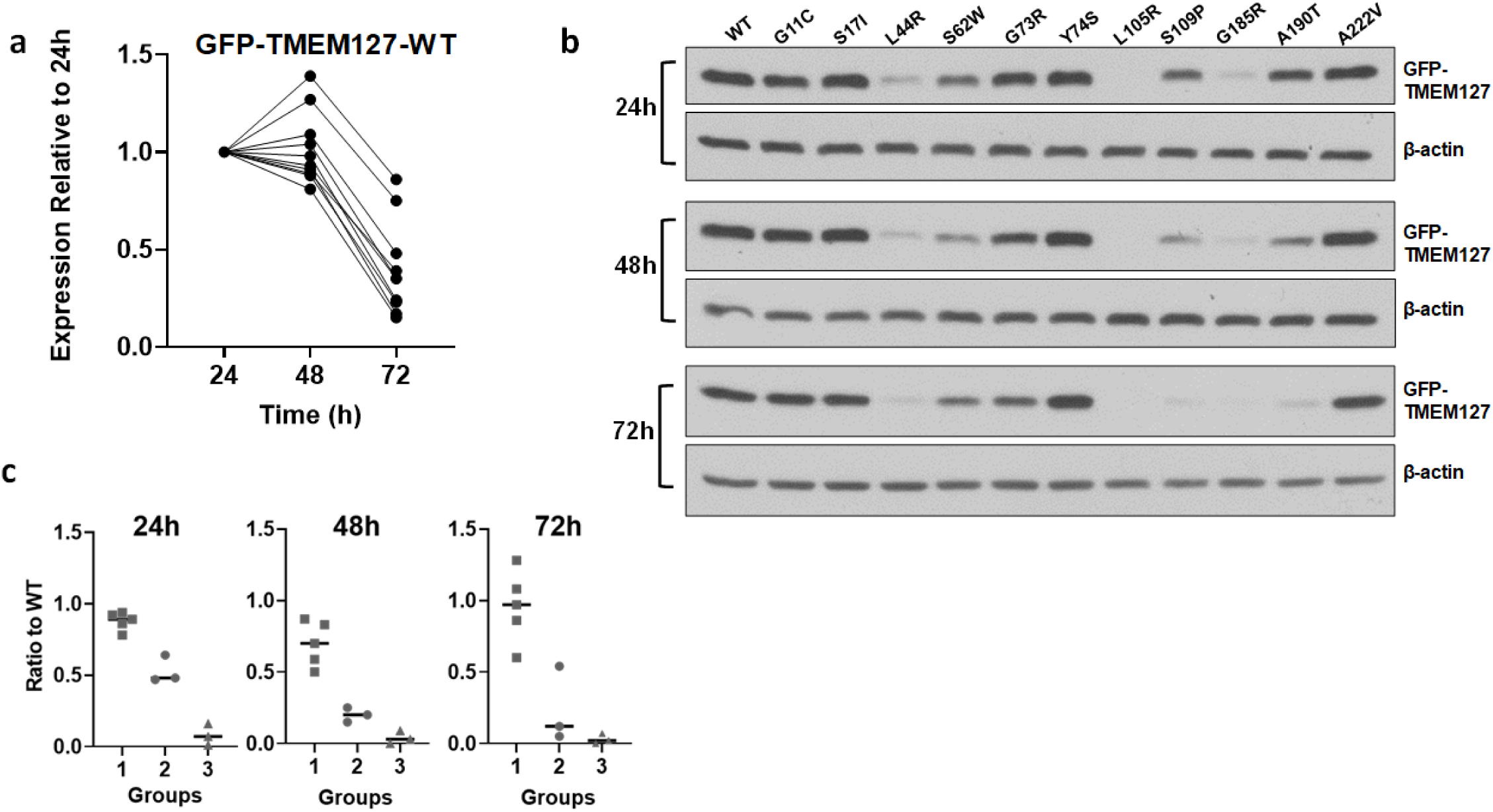
Steady-state levels of tumor-associated germline TMEM127 variants. (a) Quantification of GFP-TMEM127 wild-type (WT) abundance in immunoblots of transiently expressed HEK293FT TMEM127-KO cells 24, 48 and 72h after transfection, probed with a monoclonal GFP antibody, normalized by loading control (beta-actin) and quantified by ImageJ. Data from 11 replicate experiments are shown with first time point (24h) set to 1; (b) Representative immunoblots of HEK293FT TMEM127-KO cells transiently expressing GFP-tagged versions of the indicated constructs. The steady-state levels of WT and the various TMEM127 variants are shown at 24 hours, 48 hours, and 72 hours after transfection; (c) Quantification of immunoblots showing the expression levels of GFP-TMEM127 variants in HEK293FT-TMEM127 KO cells after transient transfection for the indicated time points. The GFP-TMEM127 band was normalized to loading control and then to WT construct in each blot. Three groups of variants were identified: 1) variants showing a pattern similar to, or slightly decreased, relative to WT; 2) variants that had at least two time points significantly decreased compared to WT; and 3) variants with minimal to no detection at all time points. All diffuse variants belonged to group 3. Most punctate variants and the single plasma membrane variant were in group 1; three punctate variants (S62W, S109P and A190T) had lower abundance than WT. Each blot was run at least twice, mostly three or more times. The three expression groups are significantly different(p<0.05) by one-way ANOVA.

**Fig. 3.**
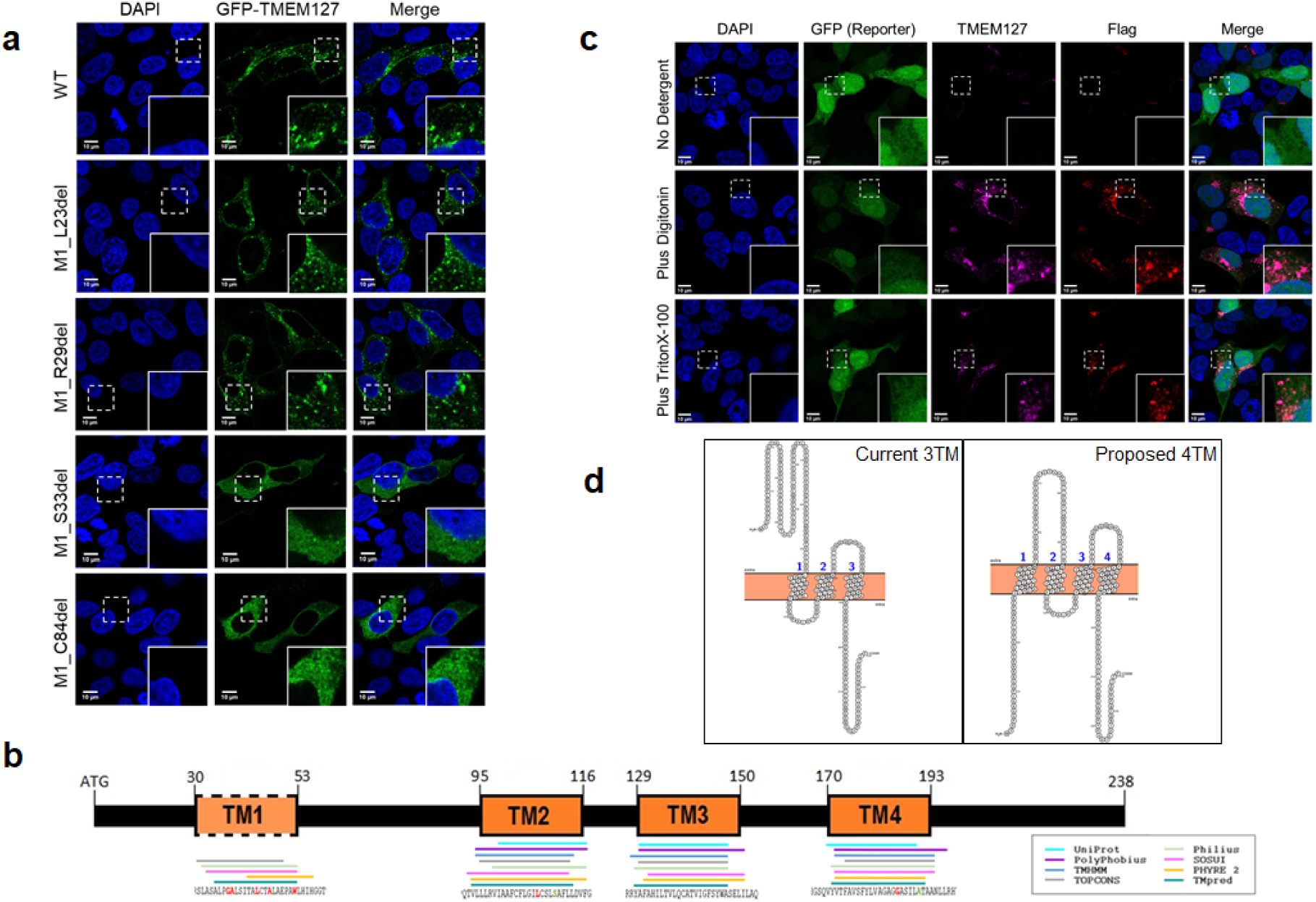
Membrane topology of TMEM127. (a) Immunofluorescence analysis of HEK293FT TMEM127-KO cells transiently expressing GFP-tagged TMEM127 versions of the indicated constructs designed to contain N-terminal truncations of 23, 29, 33 or 84 amino acids. Constructs lacking the first 23 or 29 amino acids produce punctate/endomembrane proteins similar to WT whereas truncation of 33 and 84 amino acids produce diffuse/cytoplasmic proteins suggesting that an additional transmembrane domain likely starts between residues 29 and 33. DAPI (blue) indicates nuclei; (b) Graphic representation of results from *in silico* analysis using eight distinct membrane topology prediction programs showing the predicted locations of transmembrane domains across the 238-residue TMEM127 protein. Amino acids representing ‘consensus’ delimitation of the four transmembrane (TM) domains based on patient-derived mutation and deletion mapping are outlined; (c) Immunofluorescence analysis of HEK293FT TMEM127-KO cells transiently expressing MSCV-TMEM127-Flag (tag on the C-terminus of the protein) were used for antibody accessibility assays. GFP is expressed bicistronically from this construct and identifies transfected cells. DAPI (blue) indicates nuclei. A polyclonal TMEM127 antibody recognizing the first 50 amino acids of the TMEM127 sequence was used to probe the N-terminus (magenta); a Flag monoclonal antibody was used to recognize the C-terminus (red). Top panel: neither the N-nor the C- (Flag, red) terminus of TMEM127 were detected under unpermeabilized conditions; middle panel: both N and C terminals were detected under selective permeabilization with digitonin; bottom panel: both N and C terminals were also detected under fully permeabilized conditions with TritonX. These results support a model whereby both TMEM127 termini at the plasma membrane face the cytoplasm; (d) Existing (left) and proposed (right) membrane topology for TMEM127. Our experiments are consistent with a model whereby TMEM127 is a four transmembrane protein, with both N and C termini at the plasma membrane directed to the cytoplasmic side. Image generated using the Protter visualization tool based on Philius topology predictions for transmembrane domain locations.

### Variability in expression levels of germline T*MEM127* variants

We had previously observed that pathogenic TMEM127 variant proteins were undetectable by Western blot, implying potential instability and premature degradation^7,8,11,13^. We established WT construct expression over a time course of 24, 48 and 72h after transfection in HEK293FT cells that we previously engineered to lack *TMEM127*^*11*^ (Fig. 2a), and evaluated comparatively the abundance of TMEM127 variant constructs. We detected three groups of TMEM127 variants (Fig 2b, 2c). One group (group 3) contained all diffuse variants (such as p.L44R, p.L105R and p.G185R) which displayed markedly reduced abundance at all three time points compared to the WT (Fig. 2b, 2c; Supplementary Fig. S1c, d, e). This indicated that loss of membrane binding ability most likely has a profound, negative impact on the expression of these variants. As *TMEM127* mRNA levels in tumors of patients with germline *TMEM127* truncating variants are decreased, it is plausible that reduced transcription may account for the protein deficit in diffuse variants^6^.

The remaining two groups contained the eight punctate variants. These punctate variants generally separated into two categories: those with protein levels similar to WT across at least two of the three time points tested (group 1), and those with decreased levels relative to WT (group 2), suggestive of reduced synthesis and/or instability of the variant (Fig2b-f, Supplementary FigS1c-f). Interestingly, variant proteins with steady-state levels broadly similar to WT (p.S17I, p.G73R, p.Y74S, p.A222V) were all located outside of transmembrane domains, whereas two of the three variant proteins with lower levels than WT (p.S109P, p.A190T), were both located within transmembrane domains (Fig. 2b). This observation suggested that certain missense variants occurring within transmembrane domains retain some ability to associate with membrane (Supplementary Fig S1f) but may still lead to decreased abundance and/or stability at the protein level. Importantly, these observations were validated using untagged TMEM127 constructs, indicating that this was not an artifact of the GFP tag (Supplementary Fig. S1c, d, e). These data confirm that membrane association is not grossly affected in these variants, although we cannot exclude that their membrane binding ability is modified to a subtler degree, not detectable by our methods, or if these residues are subject to posttranslational modifications. A third member of group 2, p.S62W, is located outside a putative transmembrane domain. The mechanisms explaining how these punctate variants result in decreased protein levels remain to be defined.

Two other variants are worth additional mention. First, p.G11C, a component of group 1, could not be detected using a TMEM127 polyclonal antibody targeting the N-terminal region, although it was clearly observed by an antibody that recognized the GFP tag (Supplementary Fig. S1e). It is currently unknown if this residue or region is functionally relevant in some other capacity (i.e. protein-protein interactions) and/or whether it is subject to folding or other structural modification. A second variant of interest, also mbelonging to group 1, was the plasma membrane bound variant protein (p.M214Sfs98X), which had steady-state levels comparable to WT at the earlier, but not the last time point (Supplementary Fig3c) consistent with potentially distinct temporal processing and/or regulatory mechanisms.

These results suggest that changes in TMEM127 variant abundance complements subcellular distribution analysis and may flag additional states of disrupted protein function. Importantly, several of these defects would not have been identified by pathogenicity prediction algorithms (Table 1), and these variants would have not been appropriately classified.

### Evaluation of TMEM127 membrane topology

As mentioned above, we noticed that five missense variants (p.G37R, p.A38D, p.L44R, p.A47D, p.W53S), which clustered in an N-terminal region not previously known to contain a functional domain, were diffuse (Fig. 1b, Supplementary Fig S2a). Four of these variants had clinical indicators of pathogenicity (e.g. bilateral PHEOs and/or loss of heterozygosity of the WT allele)^7-9,14,19^, in support of a functional defect. Of note, when we investigated two additional *TMEM127* variants targeting the p.A38 residue^25^, we found they were punctate, indicating that the type of amino acid substitutions impacted on the subcellular localization outcome (Supplementary Fig S2a,b). Other missense variants located upstream, p.G11C, p.S17I (Fig1d) and downstream-p.S62W (Supplementary Fig S1a), p.G73R, p.Y74S-of this cluster were also punctate, similar to WT (Fig 1c, Table 1), indicating that this observation was region-specific. Furthermore, we confirmed that an N-terminal truncating variant (p.M1_C84del) spanning this region was also diffuse despite not targeting a reported transmembrane domain (Fig. 1c, 3a, Supplementary Fig. S1b). We had previously described this construct and predicted it would be the result of a variant affecting the start codon which would lead to a shift of the start site to the next in-frame methionine (p.M85)^7^. As we have previously associated the diffuse pattern with variants that disrupt transmembrane domains, these observations prompted us to consider that an additional, previously unrecognized, transmembrane domain might reside in the N-terminal region.

To investigate this possibility, we first queried the TMEM127 amino acid sequence across several membrane topology prediction algorithms. Some algorithms predicted three, while others predicted four transmembrane domains (Fig.3b, Supplementary Table S1, Supplementary Fig S2d, e). All algorithms predicted the three transmembrane domains recognized in the consensus TMEM127 entry in NCBI, NP_001180233.1 (Fig.3b). In every instance, the additional transmembrane domain was predicted to reside in the N-terminal spanning amino acids 30-57 approximately (Fig. 3b), a region that overlapped with all the diffuse missense variants that clustered in the N-terminal region and with the p.M1_C84del variant (Fig. 3a).

We then generated a series of N-terminal truncations and observed their subcellular localization. We found that deletion of 23 residues (p.M1_L23del) and 29 residues (p.M1_R29del) maintained a punctate appearance similar to WT, but that deletion of 33 residues (p.M1_S33del) resulted in a diffuse/cytoplasmic pattern similar to p.M1_C84del (Fig. 3a). Loss of membrane binding ability for p.M1_S33del supported that a transmembrane domain was disrupted by this truncation. Moreover, it indicated that the start of this transmembrane domain was likely between residues 29 and 33, in full agreement with the observations made with patient-derived variants and with the *in silico* predictions (Fig 3b, Supplementary Table S1). Hence, these results supported the existence of an additional, fourth transmembrane domain located between residues 30 and 53 of the TMEM127 sequence.

Next, to determine whether both TMEM127 ends have the same orientation, which would be consistent with an even (i.e., four) number of transmembrane domains, we performed an antibody accessibility assay. We transiently transfected HEK293FT TMEM127-KO cells with a C-terminal, Flag-tagged, TMEM127 WT construct bicistronically expressing a GFP reporter^6^, and used either a Flag antibody to recognize the C-terminus, or the aforementioned TMEM127 antibody, directed to the first 50 amino acids of the protein, to recognize the N-terminus of TMEM127. We observed that in the absence of detergent, where membranes remain non-permeabilized, exposing only extracellularly facing epitopes, neither the N-nor the C-terminal of the TMEM127-Flag construct could be detected in transfected cells (recognized by their GFP expression, Fig 3c). This indicated that when TMEM127 is at the plasma membrane, neither terminal is extracellularly oriented (Fig. 3c, top panel). On the other hand, in the presence of digitonin, a mild detergent producing selective permeabilization of the plasma membrane, as well as Triton X-100, a harsh detergent producing full permeabilization of all membranes, both terminals were detected in the cytoplasm (Fig. 3c, middle and bottom panels). These results indicated that both TMEM127 terminals are oriented in the same direction, toward the cytoplasm, supporting an even, not an odd, number of transmembrane domains.

Altogether, our observations support the existence of a novel, fourth transmembrane domain in TMEM127 as indicated in the proposed model (Fig. 3d).

### Identification of mechanisms governing TMEM127 internalization and redistribution

TMEM127 subcellular distribution under regular culture conditions indicates that the WT protein predominantly displays a punctate endomembrane distribution, but approximately 20% is detected at the plasma membrane (Fig1a)^6^. This pattern is compatible with TMEM127 being sorted to the cell surface after its biosynthesis and routed to its endosomal/lysosomal domains after internalization from the plasma membrane. This process, generally referred to as an ‘indirect trafficking route’ common to many transmembrane proteins, requires the existence of certain cytosolic signals^34^. We initially predicted that the frameshift variant which occurred downstream of the last transmembrane domain, p.M214Sfs98X, would not affect membrane binding ability as it did not disrupt any of the transmembrane domains. Indeed, the resulting protein maintained its membrane binding ability; however, it was predominantly plasma membrane bound, a pattern that had not been previously observed in other TMEM127 variants (Fig. 1d). To explain this observation, we considered two possibilities: that the frameshift, which generated an extended stop codon resulting in a protein of 312 instead of 238 residues (Supplementary Fig.S3a), caused the protein to become too bulky to internalize, or alternatively, as a more likely scenario, that an endocytic signaling motif required for internalization was lost^34^. To address this, we generated a nonsense variant (p.T201X) which truncated a few amino acids downstream of the last transmembrane domain. The resulting protein predominantly localized to the plasma membrane, similar to p.M214Sfs98X, which was consistent with TMEM127 potentially having an endocytic signaling motif in its C-terminal, which was required for its internalization (Supplementary Fig. S3a, b), without markedly altering its expression levels (Supplementary Fig3c).

We then scanned the C-terminal of TMEM127 for potential tyrosine-based motifs (YXXΦ), where X is any amino acid and Φ is a bulky, hydrophobic acid (F,I,L,M or V), and dileucine-based motifs ([D/E]XXXL[L/I], where there are two adjacent leucines, or a leucine-isoleucine pair, and an acidic residue 4 amino acids upstream of the first leucine ^34-36^. Although these are considered to be the two classical motifs, as signaling motifs can be diverse, we also considered other less common variations such as extended-acidic, dileucine-based, endocytic motifs^37-41^. We identified two potential candidate regions: a classical tyrosine-based motif downstream of the frameshift (p.YEVI224_227), and an extended-acidic dileucine-based motif a few residues upstream of the frameshift (p.EEEEQALELL202_211). Although the putative tyrosine-based motif appeared to be the likely candidate, neither mutation of the presumed critical residues (p.Y224A, I227A) nor an in-frame deletion (p.YEVI224_227del) led to increased plasma membrane localization indicating that this region was not critical for internalization (Supplementary Fig. S3d).

In contrast, mutation of the presumed critical residues (p.L210A, p.L211A as well as p.LL210_211AA) of the putative dileucine-based motif caused significant plasma membrane localization (Supplementary Table 2, Fig. 4a, b), suggesting that these residues were functional and critical for internalization. We noted that a cluster of acidic residues (p.EEEE202_205) was located upstream of the two leucines in positions that would follow an extended-acidic dileucine motif instead of the classical dileucine motif^41^. Therefore, to further investigate this region, we conducted alanine-scanning mutagenesis spanning residues p.E202 to p.E216. We determined that two acidic residues (p.E204, p.E205) upstream of the dileucines play a major role in internalization, as mutation of either residue (p.E204A, p.E205A, as well as the double-mutant p.EE204_205AA) resulted in significant plasma membrane localization (Supplementary Table 2, Supplementary Fig. S4a, b). In addition, acidic residues located downstream of the dileucine motifs have been found to contribute to internalization efficiency in some instances, especially when a classical motif is not present^37,41^. Similarly, we determined that two residues (p.M214, p.E215) downstream of the dileucines are also required for internalization and resulted in significant plasma membrane localization when mutated (Supplementary Fig. S4c, d). Importantly, both the p.M214 and p.E215 residues are disrupted by the p.M214Sfs98X frameshift, which likely explains its plasma membrane localization. We further demonstrated the contribution of this region downstream of the dileucines to internalization by using nonsense variants. While p.E213X resulted in significant plasma membrane localization, p.E216X maintained a punctate appearance similar to WT (Supplementary Fig. S4e, f). Altogether, our data suggest that TMEM127 has an atypical, extended-acidic dileucine-based endocytic motif (EEXXXXLLXXME, where X is not a critical residue for internalization).

**Fig. 4.**
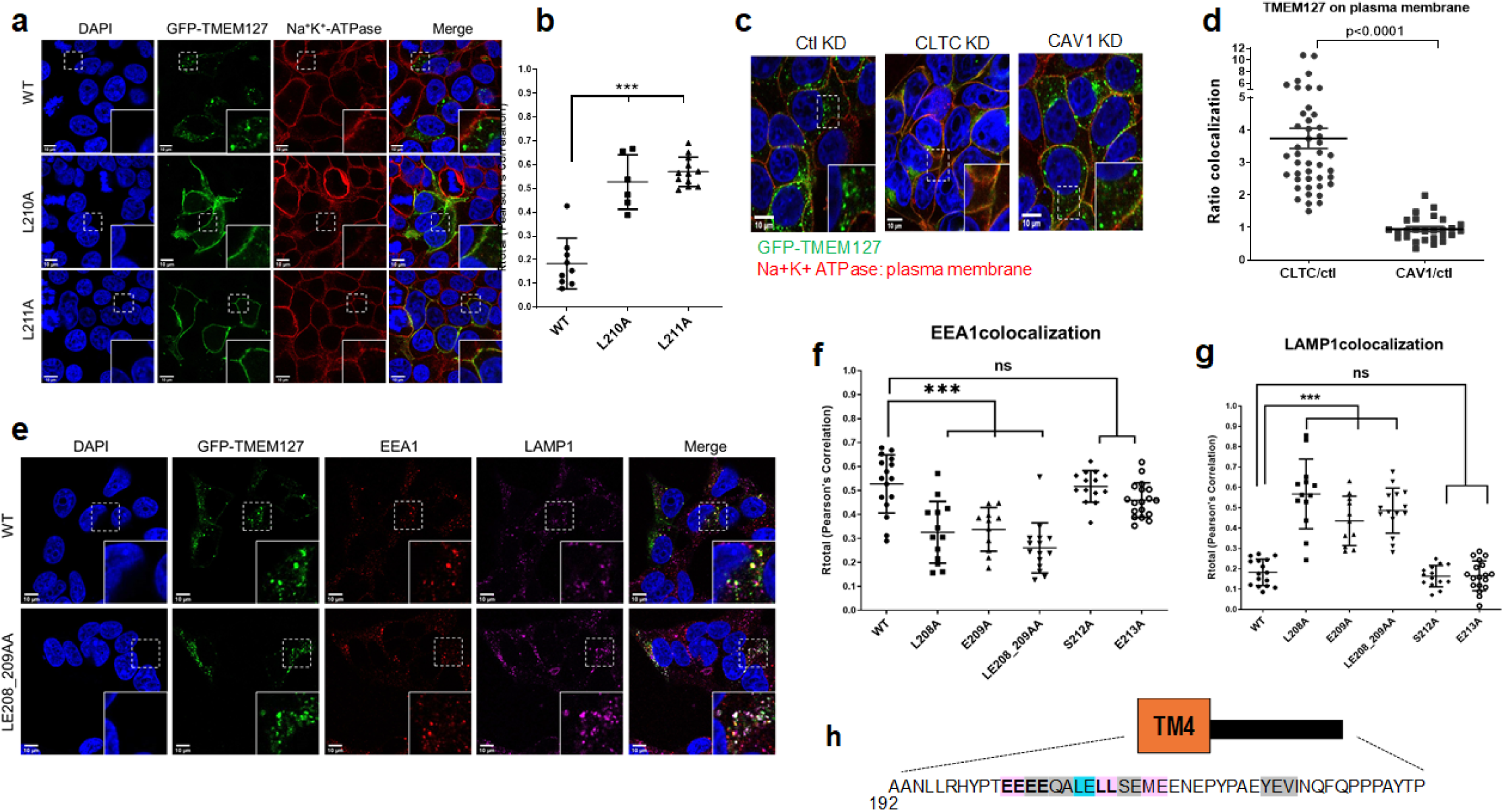
Mechanisms of internalization of TMEM127. (a) Immunofluorescence analysis of HEK293FT TMEM127-KO cells transiently expressing GFP-tagged TMEM127 with the indicated constructs. Representative confocal images of colocalization of TMEM127 (green) with Na^+^K^+^ATPase (red), a plasma membrane marker showing that mutation of both p.L210 and p.L211 result in significant plasma membrane localization of TMEM127; (b) Quantification based on Pearson’s correlation coefficient of TMEM127 and NA^+^K^+^ATPase colocalization from three independent experiments. Average and SEM are shown. Student’s t test was used for statistical calculations; p<0.001=***; (c) Representative immunofluorescence image of HEK293FT cells stably expressing GFP-tagged TMEM127 WT transduced with either scrambled shRNA (Ctl KD), shRNA against clathrin heavy chain (CLTC KD) and, from siRNA against caveolin (CAV1 KD), stained for NA^+^K^+^ATPase (red); KD= knockdown. Not shown: scrambled siRNA used as control for the CAV1 KD experiments, similar to Ctl KD shRNA; representative images shown in Supplementary Figures 5b and 5d; (d) Quantification based on Pearson’s correlation coefficient of TMEM127 and NA^+^K^+^ATPase colocalization from three independent experiments, normalized to respective scrambled shRNA or siRNA controls. Average and SEM are shown. Student’s t test was used for statistical calculations; p<0.0001=***; (e) Confocal images of HEK293FT TMEM127-KO cells transfected with either GFP-tagged TMEM127 WT or p.LE208_209AA variant (green) and probed for EEA1, an early endosomal marker (red) and LAMP1, a lysosomal marker (magenta); (f, g) Quantification of colocalization of GFP-TMEM127 WT or variants and (f) EEA1 and (g) LAMP1, based on Pearson’s correlation coefficient. Average from three independent replicates and SEM are shown. Student’s t test was used for statistical calculations; p<0.001=***; (h) Graphic summary of functionally relevant residues involved in TMEM127 internalization (pink) and redistribution(cyan), and those not critical for internalization (gray) based on mutagenesis and confocal microscopy of individual mutants. Residues 194-238 are downstream of the fourth and most distal transmembrane domain.

Transmembrane proteins containing these types of endocytic motifs in their cytoplasmic tails frequently utilize clathrin-mediated endocytosis (CME) for internalization before getting routed to their intended intracellular destination^42^. However, clathrin-independent (i.e. caveolae/lipid raft dependent) endocytosis mechanisms also exist and, in some cases, proteins can utilize both mechanisms^43^. To further define the mechanism governing the internalization of TMEM127, we generated HEK293FT cells stably expressing GFP-tagged TMEM127 WT and, using RNA interference, targeted two major proteins, clathrin heavy chain (CLTC) and caveolin 1 (CAV1), involved in their respective pathways. We obtained efficient knockdown for both targets (Supplementary Fig.S5a,5b) and observed that upon knockdown of CLTC, but not CAV1, TMEM127 WT accumulated and became localized predominantly to the plasma membrane (Fig. 4c, d; Supplementary Fig.S5c, 5d). This suggested that internalization of TMEM127 was blocked by clathrin deficiency, in agreement with TMEM127 utilizing CME, but likely not caveolin-dependent endocytosis, for internalization.

Lastly, we noticed that mutation of several residues outside of those required for internalization resulted in an apparent intracellular redistribution under the regular culture conditions of our experiments (Table 2). In particular, mutation of two residues directly upstream of the dileucines, p.L208 and p.E209, led to significantly higher colocalization with the lysosomal marker LAMP1 and concomitantly lower colocalization with the early endosomal marker EEA1 when compared to the WT and other surrounding punctate variant proteins (Fig. 4e, f, g; Supplementary Fig. S5e). We also noted that p.L208A and p.E209A (as well as the double-mutant p.LE208_209AA) were more likely to have enlarged ring-like vesicles (Supplementary Fig. S5f), while displaying abundance levels broadly similar to WT by immunoblots (Supplementary Fig. S5g). This observation suggests that additional residues surrounding the endocytic motif, specifically p.L208 and p.E209, but potentially other residues as well, appear to be necessary for efficient subvesicular distribution.

## DISCUSSION

In this study, we functionally characterized 21 tumor-associated, germline *TMEM127* variants by evaluating their subcellular localization and steady-state levels. By combining these results, we determined that 15 of these 21 variants, including 9 of 15 missense variants, can be classified as pathogenic or likely pathogenic, through loss of membrane binding ability, stability or internalization capability. As involvement in the endo-lysosomal system appears to be relevant to the function of TMEM127^10,15^, we considered any variant that produced a diffuse/cytoplasmic distribution to be highly disruptive and thus pathogenic (group 3). Likewise, a plasma membrane bound variant that is defective in internalization is likely pathogenic. We also considered variants which affect protein stability to be likely pathogenic (group 2) although additional work is needed to characterize the exact deficit leading to this instability. In addition, we propose that the six missense variants that were punctate/endomembrane and relatively stable remain classified as variants of uncertain significance (VUS) until additional functional assays are developed.

Furthermore, our observations with several of these patient-derived variants revealed the presence of two novel structure-function features of TMEM127. First, we established that an additional transmembrane domain resides in the N-terminal indicating that TMEM127 is a four-, not a three-, transmembrane domain protein. Four transmembrane domain proteins encompass a wide array of proteins and protein families including ion channels, claudins, connexins, and tetraspanins with varying functions^44,45^. As the function of TMEM127 remains largely unknown, this newly defined topology will allow for assessment of possible features by structurally and functionally comparing TMEM127 to other four transmembrane proteins.

Second, we uncovered an atypical, extended-acidic, dileucine-based, endocytic motif in the C-terminal of TMEM127 which is required for its internalization. Although we have previously observed that TMEM127 WT dynamically localizes to the plasma membrane and, more predominantly, to endosomal and lysosomal membranes^6,10,15^, we had limited understanding of the mechanisms governing this distribution. Here we identified critical residues required for TMEM127 internalization from the plasma membrane, EEXXXXLLXXME, and that this process is clathrin-mediated. Although not extensively described in the literature, atypical dileucine motifs have been found in other transmembrane proteins, including several endo-lysosomal membrane proteins and can be as effective for internalization as the classical dileucine motif^37,39,40^. Notably, the efficacy of this endocytic signal would be absent in all TMEM127 variants that result in a loss of membrane binding ability (a distinguishing feature of several missense, indel, frameshift and truncating variants) or become plasma membrane bound. Additionally, the integrity of residues surrounding the dileucines, specifically p.L208 and p.E209, appears to be required for proper intracellular positioning along endosomal compartments, suggesting they contribute to proper temporo-spatial localization of TMEM127. Additional studies will be necessary to precisely define whether these residues are involved in sorting and/or trafficking.

Overall, our findings have prompted us to propose a revised TMEM127 protein model which incorporates these novel structure-function features (Fig. 2d, Fig. 3h). While this model can serve as a guide in evaluating the pathogenicity of a variant, *in vitro* functional assessments are still required for uncharacterized missense variants even if they affect the same residue as a variant already assessed. Pathogenic variants that occur within and disrupt transmembrane domains are highly likely to result in disrupted TMEM127 function. Currently, we recommend that clinical information including classic features associated with heritability (e.g., tumor multiplicity, positive family history with variant segregation with the phenotype) as well as evidence of loss of heterozygosity of the WT allele be evaluated before ascribing pathogenicity.

In summary, we have expanded our ability to predict the pathogenicity of *TMEM127* variants by identifying novel domains and residues required for normal protein function. These findings will assist in the clinical interpretation of genetic screening results. As more variants are identified and/or as the physiological role of TMEM127 becomes clearer, additional screening methods will be developed for future functional assessments.

## SUPPLEMENTARY INFORMATION

The online version of this article contains supplementary material, which is available to authorized users.

## ACKNOWLEDGEMENTS

We thank Dr. Lois Mulligan and her lab at Queens University, Kingston, Ontario, Canada for providing the short hairpin RNAs (shRNAs) against the clathrin heavy chain (CLTC) and for critical review of this manuscript, and to our collaborators who shared their variant data. Confocal images were generated in the Optical Imaging Facility, which is supported by UT Health San Antonio, NIH-NCI P30 CA54174 (CTRC at UTHSCSA) and NIH-NIA P01A.

