## Supplementary Data for "Functional characterization of germline variants in the *TMEM127* tumor suppressor reveals novel insights into its membrane topology and trafficking"

Supplementary Figure S1
Supplementary Figure S2
Supplementary Table S1
Supplementary Figure S3
Supplementary Table S2
Supplementary Figure S4
Supplementary Figure S5

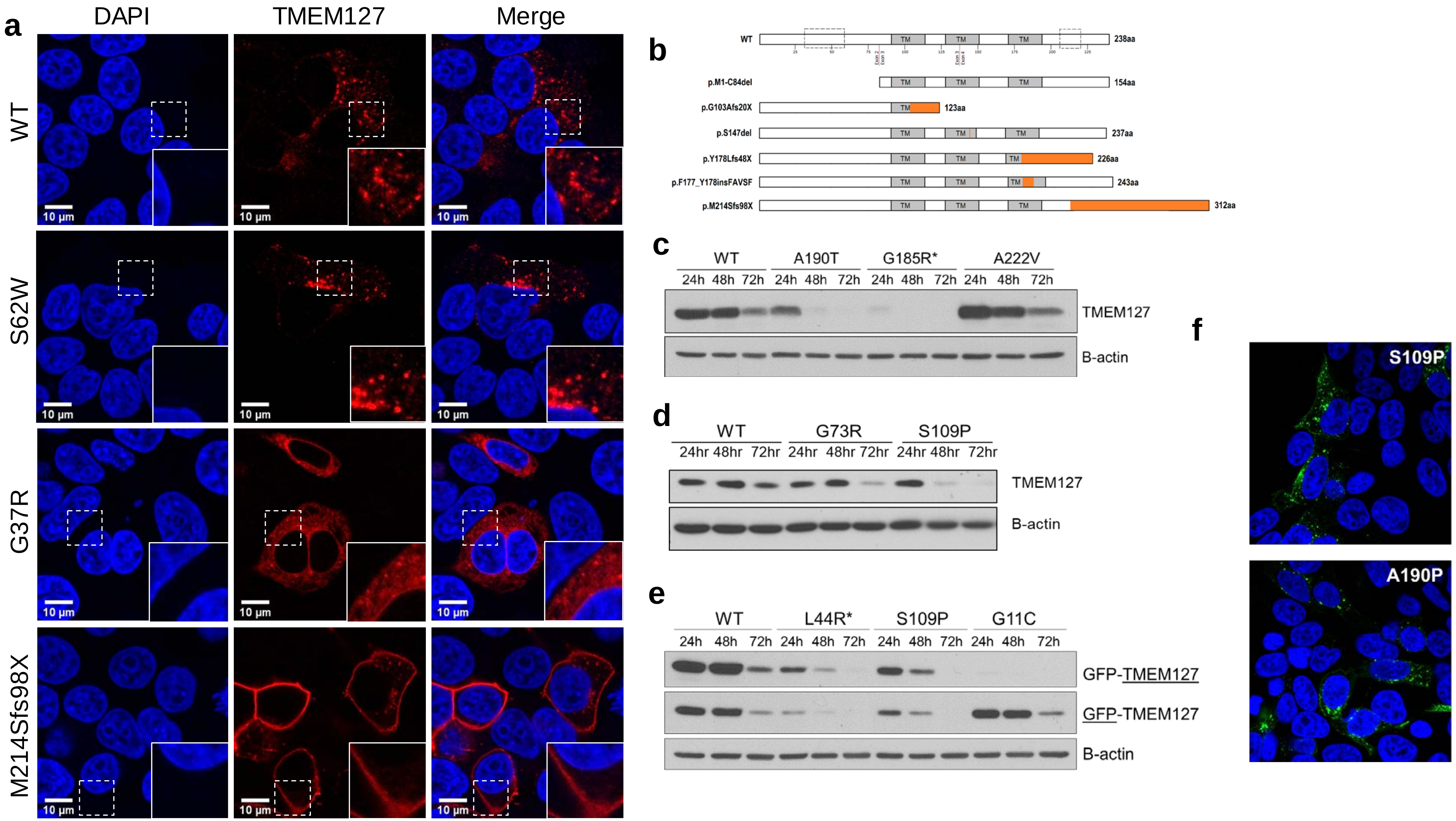
a
DAPI
TMEM127
Merge
WT
S62W
G37R
M214Sfs98X
10 µm
b
WT
TM
238aa
p.M1-C84del
154aa
p.G103Afs20X
123aa
p.S147del
237aa
p.Y178Lfs48X
226aa
p.F177_Y178insFAVSF
243aa
p.M214Sfs98X
312aa
c
WT
A190T
G185R*
A222V
24h 48h 72h
TMEM127
B-actin
d
WT
G73R
S109P
24hr 48hr 72hr
TMEM127
B-actin
e
WT
L44R*
S109P
G11C
24h 48h 72h
GFP-TMEM127
GFP-TMEM127
B-actin
f
S109P
A190P

Supplementary Fig. S1.

(a) Subcellular localization of untagged TMEM127 variant proteins transiently transfected in HEK293FT TMEM127-KO cells. Representative variants from each subcellular localization pattern observed were generated using an untagged pCMV6-TMEM127 construct and transfected into HEK293FT TMEM127-KO cells. Results were concordant with the corresponding GFP-TMEM127 variant protein.

(b) Graphic depiction of the effect of frameshift, truncating, insertion and deletion variants on the TMEM127 protein. These variant proteins were all diffuse except for p.M214Sfs98X which was plasma membrane bound. Changes from the WT protein sequence are represented in orange. The N- and C- regions of interest to investigate for an additional transmembrane and an endocytic motif, respectively, are marked by dashed rectangles.

(c, d) Untagged TMEM127 variant proteins transiently expressed in HEK293FT TMEM127-KO cells had similar steady-state level outcomes as their corresponding GFP-tagged TMEM127 variant proteins. This indicated that the GFP moiety is not influencing the localization of TMEM127..

(e) GFP-tagged TMEM127 variant proteins transiently expressed in HEK293FT TMEM127-KO cells probed for GFP and TMEM127. The variant p.G11C could only be detected with the GFP antibody, not the TMEM127 antibody which targets an N-terminal region in the first 40 amino acids (Bethyl Labs). Although the subcellular localization and steady state levels of p.G11C are similar to WT, if this residue or region is functional in some other capacity (i.e. protein-protein interactions), this observation could support pathogenicity.

(f) Immunofluorescence of the only two missense variants (p.S109P and p.A190T) localized to transmembrane domains that retain punctate appearance. However, both variants show lower steady state levels by immunoblot, compared to the WT [see (c)  and (d)]

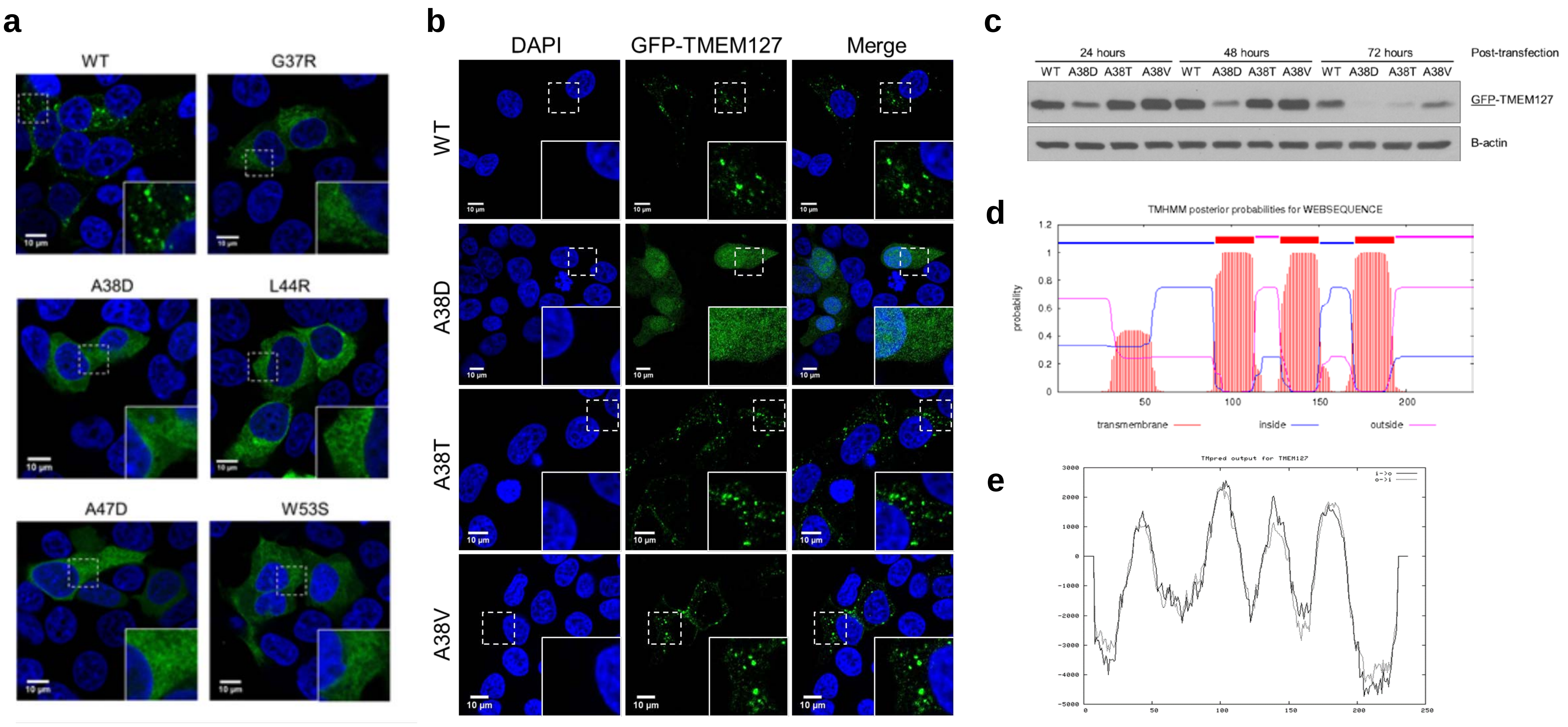
a
WT
G37R
A38D
L44R
A47D
W53S
10 μm
b
DAPI
GFP-TMEM127
Merge
WT
A38D
A38T
A38V
10 μm
c
24 hours
48 hours
72 hours
Post-transfection
WT A38D A38T A38V WT A38D A38T A38V WT A38D A38T A38V
GFP-TMEM127
B-actin
d
TMHMM posterior probabilities for WEBSEQUENCE
probability
transmembrane
inside
outside
e
TMpred output for TMEM127
i->o
o->i

Supplementary Fig. S2

(a) Subcellular localization of GFP-TMEM127 variant proteins transiently transfected in HEK293FT TMEM127-KO cells. Unlike the punctate/endomembrane appearance of the WT, five tumor-associated germline TMEM127 missense variants (p.G37R, p.A38D, p.L44R, p.A47D, p.W53S), clustered at an N-terminal region not previously associated with a functional domain, produced diffuse/cytoplasmic proteins, a pattern that was previously observed in variants that disrupt one or more transmembrane domains. Residues upstream (p.S17I, see Fig1a) and downstream (p.S62W, see Suppl. Fig1a) display a punctate/ endomembrane pattern, suggesting that this profile is specific to this region of the TMEM127 sequence.

(b) Subcellular localization of GFP-TMEM127 variants targeting alanine 38 in HEK293FT TMEM127-KO cells. Although the subcellular localization of p.A38D, resulting from a patient-derived germline TMEM127 variant, was diffuse/cytoplasmic, both p.A38T and p.A38V, resulting from potentially germline and somatic *TMEM127* variants, retained a punctate, endomembrane appearance similar to WT. These results indicate that amino acid substitutions on the same residue can produce different subcellular localization patterns.

(c) Immunoblot analysis of the constructs shown in (b) 24, 48 and 72h after transfection. In agreement with the confocal microscopy analysis, the steady-state levels of p.A38D were markedly lower than those of WT, while those of  p.A38T and p.A38D were generally similar to WT.

(d, e) Graphical images from the membrane topology prediction programs (c) TMHMM and (d) TMpred predicting three and four transmembrane domains for TMEM127, respectively. All programs utilized predicted the three previously reported transmembrane domains but had discrepant results for the additional N-terminal transmembrane domain. Although a potential domain in the N-terminal region was detected by the TMHMM program, it did not cross the threshold for calling.

Supplementary Table S1. Summary of Predictions from Various Protein Prediction Programs for TMEM127 Membrane Topology

| **Protein Topology Prediction Tool** | **Number of Predicted TM Domains** | **Location of Predicted TM Domains** | **N-Terminal/C-Terminal Orientation Prediction** |
|---|---|---|---|
| UniProt | 3 | 96-116, 130-150, 169-189 | N/A |
| PolyPhobius | 3 | 91-116, 130-154, 171-196 | SignalPeptide-Outside/Inside |
| TMHMM | 3 | 91-113, 128-150, 171-193 | Inside/Outside |
| TOPCONS | 4 | 30-50, 92-112, 130-150, 173-193 | Inside/Inside |
| Philius | 4 | 31-53, 95-116, 129-150, 170-193 | Cytoplasmic/Cytoplasmic |
| SOSUI | 4 | 32-53, 89-111, 132-154, 171-193 | N/A |
| PHYRE 2 | 4 | 42-57, 90-116, 131-154, 171-191 | Extracellular/Extracellular |
| TMpred | 4 | 34-53, 90-113, 129-150, 169-193 | N-terminus Inside |

TM = Transmembrane, N/A = Not Available; UniProt (www.uniprot.org);
PolyPhobius (www.phobius.sbc.su.se/poly.html )
TMHMM (www.cbs.dtu.dk/services/TMHMM/ )
TOPCONS (http://topcons.cbr.su.se/pred/ )
Philius (www.yeastrc.org/philius/pages/philius/runPhilius.jsp )
SOSUI (http://harrier.nagahama-i-bio.ac.jp/sosui/ )
PHYRE 2 (www.sbg.bio.ic.ac.uk/phyre2/ )
TMPred (https://embnet.vital-it.ch/software/TMPRED_form.html )

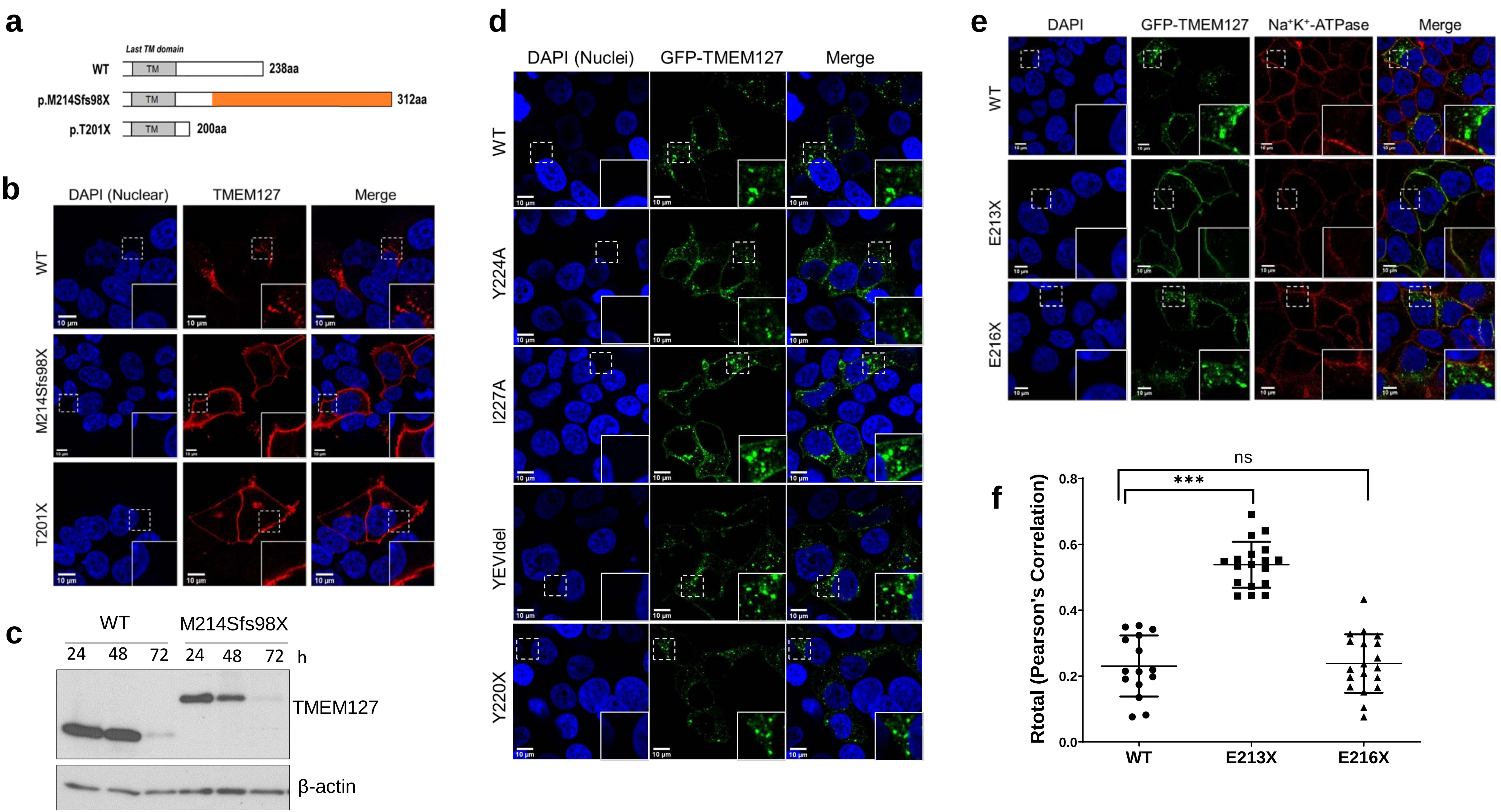
a
Last TM domain
WT
TM
238aa
p.M214Sfs98X
312aa
p.T201X
200aa
b
DAPI (Nuclear)
TMEM127
Merge
WT
M214Sfs98X
T201X
10 µm
c
WT
M214Sfs98X
24 48 72 24 48 72 h
TMEM127
β-actin
d
DAPI (Nuclei)
GFP-TMEM127
Merge
WT
Y224A
I227A
YEVIdel
Y220X
e
DAPI
GFP-TMEM127
Na+K+-ATPase
Merge
WT
E213X
E216X
f
Rtotal (Pearson's Correlation)
0.0
0.2
0.4
0.6
0.8
ns
***
WT
E213X
E216X

Supplementary Fig. S3

(a) Diagram of the C-terminal of TMEM127 encompassing the most distal transmembrane domain (TM), displaying WT, and two additional variants, p.M214Sfs98X, a patient-derived, tumor-associated, stop loss variant that caused a frameshift and extended open reading frame (312 amino acids), and p.T201X, generated to further evaluate this region.

(b) Subcellular localization of GFP-TMEM127 variant proteins shown in (a) transiently transfected in HEK293FT TMEM127-KO cells. Truncation after the last transmembrane domain (p.T201X) resulted in plasma membrane localization of the protein, similar to the patient-derived TMEM127 variant p.M214Sfs98X, in agreement with an internalization motif downstream of the distal transmembrane domain.

(c) Immunoblot showing the steady-state levels of the plasma membrane-bound variant, p.M214Sfs98X over three time points. This variant had a faster decrease in its steady-state levels compared to WT at the last time point (72h), but similar to WT at earlier time points (24 and 48h). B-actin is used as a loading control.

(d) Immunofluorescence analysis of mutation of putative critical residues (p.Y224A, p.I227A), in-frame deletion of the potential motif (p.YEVI224_227del) and truncation preceding the potential motif (p.Y220X) had no effect on the subcellular localization of the protein. All mutants displayed a punctate distribution similar to WT, suggesting that internalization of TMEM127 is not dependent on a putative canonical tyrosine-based endocytic motif, YXXΦ.

(e) Immunofluorescence analysis of truncating mutants E213X and E216X co-stained with the plasma membrane marker $Na^+K^+$ATPase show plasma membrane distribution, indicating that residues downstream of 213 but upstream of 216 are required for TMEM127 internalization.

(f) Quantification of the colocalization shown in (e) based on Pearson's correlation coefficient of TMEM127 and $Na^+K^+$ATPase colocalization from three independent experiments. Average and SEM are shown. Student's t test was used for statistical calculations; $p<0.001$=***

Supplementary Table S2. C-Terminal Truncating and Alanine Scanning TMEM127 Variants and Their Effect on the Plasma Membrane Localization of TMEM127

| Construct Name | 201 | 202 | 203 | 204 | 205 | 206 | 207 | 208 | 209 | 210 | 211 | 212 | 213 | 214 | 215 | 216 | Plasma Membrane |
|---|---|---|---|---|---|---|---|---|---|---|---|---|---|---|---|---|---|
| WT | T | E | E | E | E | Q | A | L | E | L | L | S | E | M | E | E | ++ |
| p.EEEE202_205AAAA | T | ***A*** | ***A*** | ***A*** | ***A*** | Q | A | L | E | L | L | S | E | M | E | E | ++++++ |
| p.EE202_203AA | T | ***A*** | ***A*** | E | E | Q | A | L | E | L | L | S | E | M | E | E | +++ |
| p.EE204_205AA | T | E | E | ***A*** | ***A*** | Q | A | L | E | L | L | S | E | M | E | E | ++++++ |
| p.E204A | T | E | E | ***A*** | E | Q | A | L | E | L | L | S | E | M | E | E | +++++ |
| p.E205A | T | E | E | E | ***A*** | Q | A | L | E | L | L | S | E | M | E | E | +++++ |
| p.Q206A* | T | E | E | E | E | ***A*** | A | L | E | L | L | S | E | M | E | E | + |
| p.LE208_209AA* | T | E | E | E | E | Q | A | ***A*** | ***A*** | L | L | S | E | M | E | E | + |
| p.L208A* | T | E | E | E | E | Q | A | ***A*** | E | L | L | S | E | M | E | E | + |
| p.E209A* | T | E | E | E | E | Q | A | L | ***A*** | L | L | S | E | M | E | E | + |
| p.LL210_211AA | T | E | E | E | E | Q | A | L | E | ***A*** | ***A*** | S | E | M | E | E | ++++++ |
| p.L210A | T | E | E | E | E | Q | A | L | E | ***A*** | L | S | E | M | E | E | +++++ |
| p.L211A | T | E | E | E | E | Q | A | L | E | L | ***A*** | S | E | M | E | E | ++++++ |
| p.S212A* | T | E | E | E | E | Q | A | L | E | L | L | ***A*** | E | M | E | E | + |
| p.E213A | T | E | E | E | E | Q | A | L | E | L | L | S | ***A*** | M | E | E | ++ |
| p.E213X | T | E | E | E | E | Q | A | L | E | L | L | S | ***X*** | | | | ++++++ |
| p.M214A | T | E | E | E | E | Q | A | L | E | L | L | S | E | ***A*** | E | E | +++++ |
| p.M214Sfs98X** | T | E | E | E | E | Q | A | L | E | L | L | S | E | ***S*** | ***C*** | ***S*** | ++++++ |
| p.E215A | T | E | E | E | E | Q | A | L | E | L | L | S | E | M | ***A*** | E | ++++ |
| p.E216A | T | E | E | E | E | Q | A | L | E | L | L | S | E | M | E | ***A*** | ++ |
| p.E216X | T | E | E | E | E | Q | A | L | E | L | L | S | E | M | E | ***X*** | ++ |
| CRITICAL RESIDUES FOR TMEM127 INTERNALIZATION | | | | **E** | **E** | **X** | **X** | **X** | **X** | **L** | **L** | **X** | **X** | **M** | **E** | | |

'+' Represents approximately 10% of signal (based on Pearson's correlation coefficient calculated using ImageJ). If ≥40% signal observed co-localizing with the plasma membrane marker, the mutated residue was considered to be a critical residue for internalization. * These LE residues (underlined in the consensus motif) do not appear to be critical for internalization but may be important for intracellular vesicular redistribution, trafficking and/or processing. ** Patient-derived mutation. Mutated residues are shown in blue.

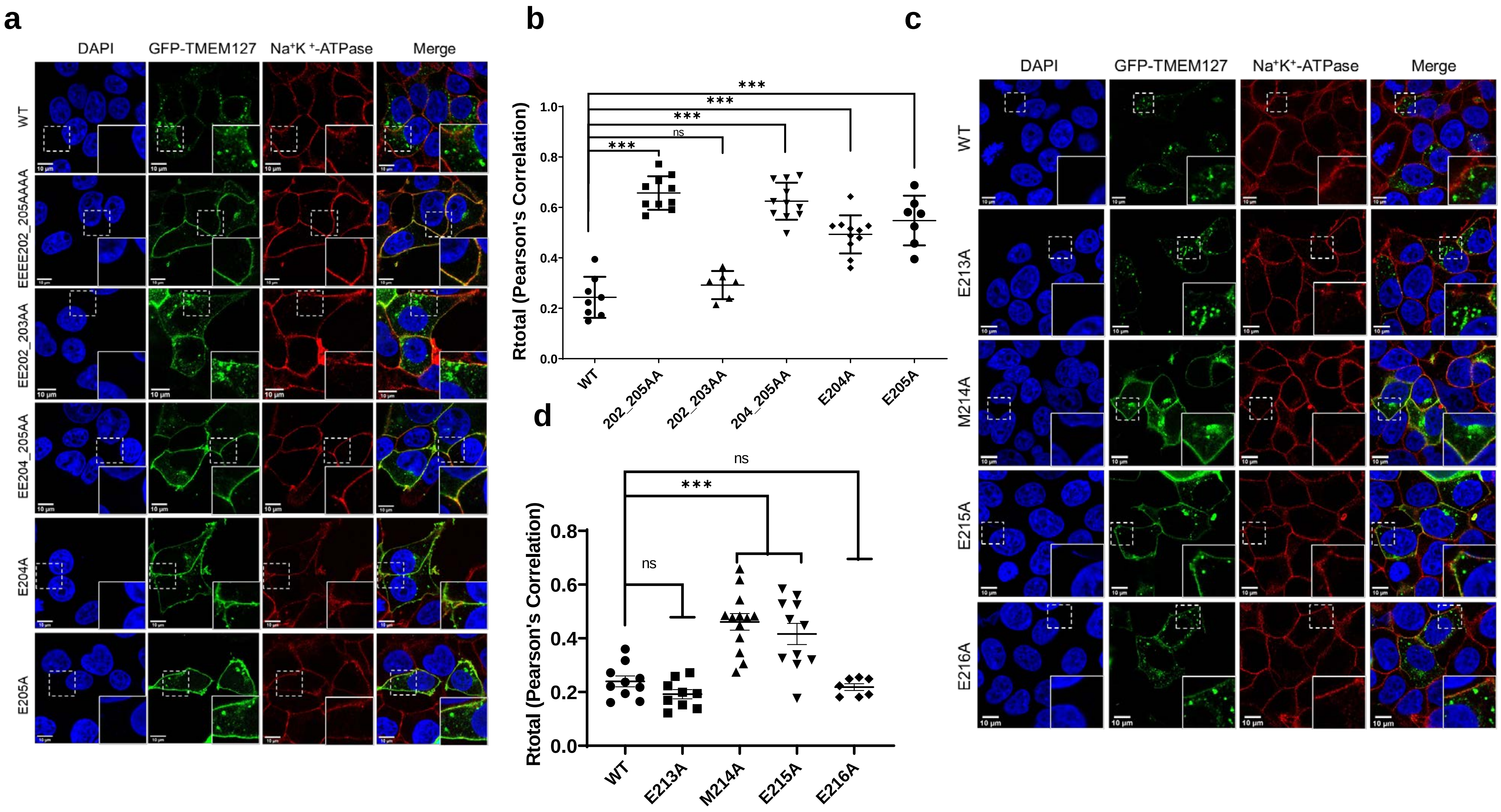

a
DAPI
GFP-TMEM127
Na⁺K⁺-ATPase
Merge
WT
EEEE202_205AAAA
EE202_203AA
EE204_205AA
E204A
E205A
b
Rtotal (Pearson's Correlation)
1.0
0.8
0.6
0.4
0.2
0.0
***
***
***
ns
***
WT
202_205AA
202_203AA
204_205AA
E204A
E205A
c
DAPI
GFP-TMEM127
Na⁺K⁺-ATPase
Merge
WT
E213A
M214A
E215A
E216A
d
Rtotal (Pearson's Correlation)
0.8
0.6
0.4
0.2
0.0
ns
***
ns
WT
E213A
M214A
E215A
E216A

Supplementary Fig. S4

(a) Immunofluorescence analysis of mutations of four acidic residues to alanines (p.EEEE202_205AAAA) co-stained with the plasma membrane marker $Na^{+}K^{+}$ATPase resulted in significant plasma membrane localization of the GFP-tagged TMEM127 protein. Subsequent mutations revealed that p.E204 and/or p.E205 were critical for internalization whereas p.E202 and p.E203 may play a minor role. These analyses indicate that E204/E205 are required for TMEM127 internalization from the plasma membrane.

(b) Quantification of the colocalization shown in (a) based on Pearson's correlation coefficient of TMEM127 and $Na^{+}K^{+}$ATPase colocalization from three independent experiments. Average and SEM are shown. Student's t test was used for statistical calculations; $p<0.001$=***

(c) Immunofluorescence analysis of mutations of p.M214 and p.E215 residues, downstream of the critical dileucines co-stained with the plasma membrane marker $Na^{+}K^{+}$ATPase. When mutated to alanines, either residue resulted in moderate plasma membrane localization of GFP-tagged TMEM127. In contrast, the adjacent residues p.E213A and p.E216A, both maintained an intracellular punctate appearance similar to WT. These results suggest that at least two residues downstream of the 210-211 dileucines, M214 and E215, also contribute to TMEM127 internalization.

(d) Quantification of the colocalization shown in (e) based on Pearson's correlation coefficient of TMEM127 and $Na^{+}K^{+}$ATPase colocalization from three independent experiments. Average and SEM are shown. Student's t test was used for statistical calculations; $p<0.001$=***

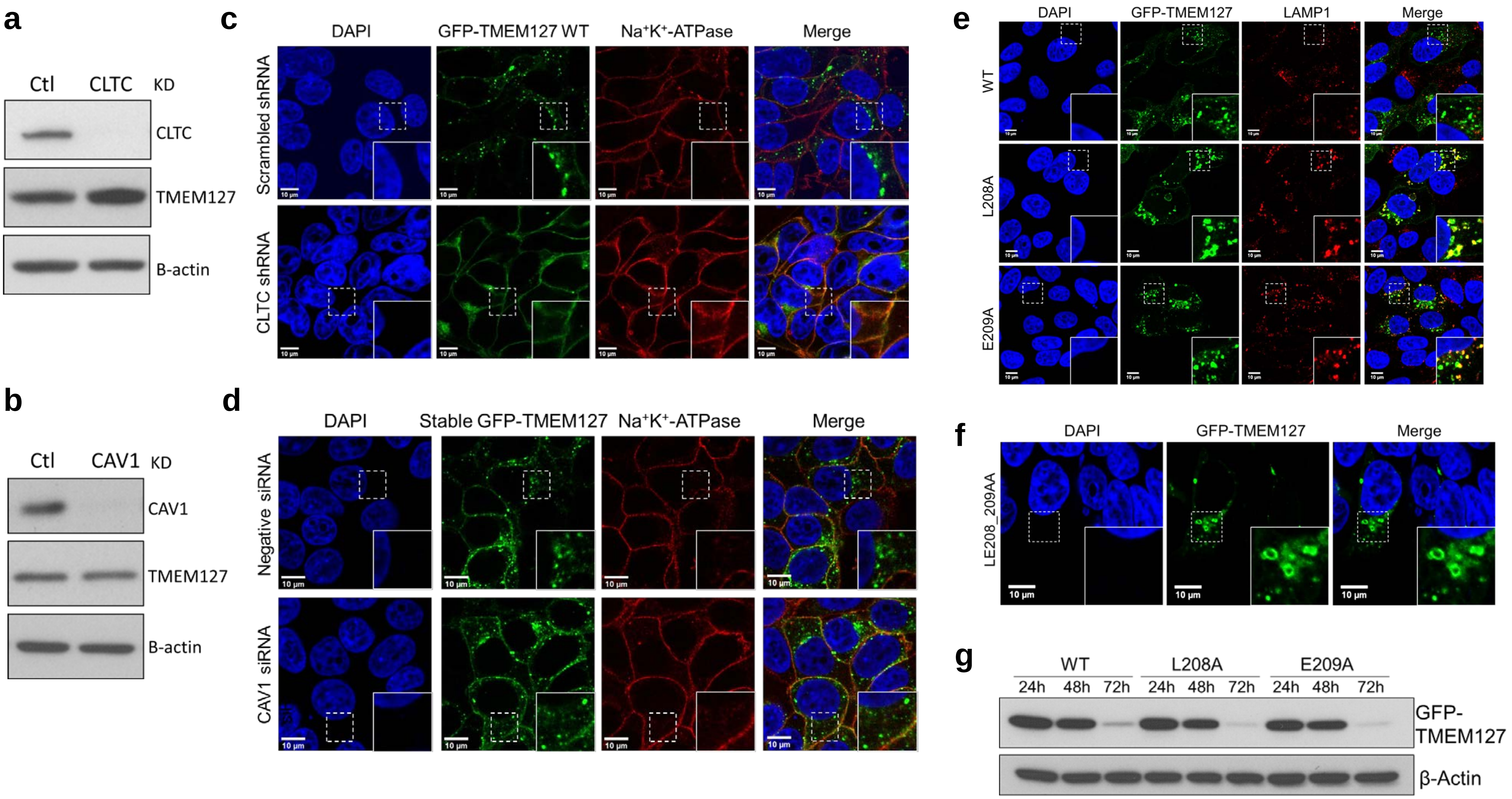

a
Ctl
CLTC
KD
CLTC
TMEM127
B-actin
b
Ctl
CAV1
KD
CAV1
TMEM127
B-actin
c
DAPI
GFP-TMEM127 WT
Na⁺K⁺-ATPase
Merge
Scrambled shRNA
CLTC shRNA
10 µm
d
DAPI
Stable GFP-TMEM127
Na⁺K⁺-ATPase
Merge
Negative siRNA
CAV1 siRNA
10 µm
e
DAPI
GFP-TMEM127
LAMP1
Merge
WT
L208A
E209A
f
DAPI
GFP-TMEM127
Merge
LE208_209AA
10 µm
g
WT
L208A
E209A
24h
48h
72h
GFP-TMEM127
β-Actin

Supplementary Fig. S5

(a) Immunoblot showing efficiency of CLTC knockdown in three independent experiments. TMEM127 abundance is increased after CLTC knockdown. B-actin is a loading control.

(b) Immunofluorescence analysis of HEK293FT cells stably expressing GFP-tagged TMEM127 WT transduced with scrambled shRNA or equal amounts of two shRNA constructs targeting clathrin heavy chain (CLTC). Images were taken 72h after transduction. Colocalization of GFP-TMEM127 signal and Na+K+-ATPase, a plasma membrane marker (red) was increased after CLTC knockdown.

(c) Immunoblot showing efficiency of CAV1 knockdown in two independent experiments. TMEM127 abundance is unchanged after CAV1 knockdown. B-actin is a loading control.

(d) Immunofluorescence analysis of HEK293FT cells stably expressing GFP-tagged TMEM127 WT treated with a control scrambled siRNA or CAV1 siRNA. No significant differences in localization of TMEM127 were observed between the two treatment groups indicating that caveolin dependent endocytosis is unlikely to serve as a major mechanism of internalization for TMEM127.

(e) Immunofluorescence analysis of HEK293FT TMEM127 KO transfected with GFP-TMEM127 constructs mutated for p.L208A, and p.E209A (two residues adjacent to the two critical dileucines) co-stained with the lysosomal membrane marker LAMP1. These mutants do not result in increased plasma membrane localization but display increased co-localization with LAMP1 compared to WT, suggesting an accumulation at the lysosome.

(f) Immunofluorescence analysis of HEK293FT TMEM127 KO with a double GFP-TMEM127 mutant construct p.LE208_209AA show enlarged enlarged vesicles

(g) Immunoblot of HEK293FT TMEM127 KO cells transiently transfected with p.L208A and p.E209A show a similar pattern to WT suggesting that the variants are not being degraded faster than WT despite their increased localization at the lysosome.
